# Molecular dynamic investigation for ROCO-4 kinase inhibitor as treatment options for parkinsonism

**DOI:** 10.1101/2023.10.21.563412

**Authors:** Kankana Datta, Lokesh Ravi

**Affiliations:** Department of Biological Sciences, University of Trieste, Italy; Department of Food Technology, Ramaiah University of Applied Sciences, Karnataka, India

**Keywords:** Parkinson Disease, LRRK2, ROCO4, GROMACS, Mongolicain-A, Bacoside-A

## Abstract

Parkinson’s disease (PD) is a neurodegenerative condition that degenerates dopaminergic neurons and is characterized by motor disabilities like rigidity, bradykinesia, postural instability, and resting tremors. Although the etiology of PD remains uncertain, familial and sporadic forms of PD are known to be predominately caused by the G2019S mutation present in the kinase domain of LRRK2. Therefore, it might be possible to treat Parkinson’s by inhibiting the kinase action of the mutated LRRK2 protein. In order to find possible inhibitors, 3069 phytochemicals were examined for their ability to bind ROCO4 kinase, which has structural and functional similarities to the LRRK2 protein. Open-source bioinformatics tools were used to determine the binding affinities of phytochemicals to the native and mutant variants of the protein. Mongolicain-A exhibited high specificity and selectivity towards the G2019S mutation of the ROCO4 protein with -12.3 Kcal/mol binding affinity, whereas Bacoside-A displayed high affinity (11.4 Kcal/mol) for the target protein, but lacked specificity towards the mutant form of the protein. Based on molecular simulation studies., RMSD, RMSF, SASA, potential energy, and hydrogen bond analysis, it was suggested that Mongolicain-A may be an effective inhibitor of the G2019S mutation and a promising drug molecule to address PD.

## Introduction

Parkinson’s disease (PD) affects 10 million individuals globally and is the second most prevalent neurological disorder^1^.The presence of Lewy bodies in dopaminergic neurons and its substantial degeneration within the substantia nigra of the midbrain are pathological hallmarks of the disease^2^ in addition to physical manifestations of bradykinesia, tremor, rigidity, sleep disorders, and constipation^3^.The etiology of PD is vague, ranging from environmental exposure to pesticides, genetic mutations, or traumatic brain injury affecting individuals between 50 and 65 years of age^1^.

However, the majority of familial and sporadic instances of PD have been found to have the pathogenic mutation G2019S in the kinase domain of Leucine Rich Repeat Kinase (LRRK2). The prevalence of this mutation ranges between 2–40% in varied populations^4,5^ and is known to inhibit neuronal differentiation and reduce levels of transcription factors like NRF21 and PITX3 that support neurogenesis and dynamic differentiation of dopaminergic neurons^6,7^. Additionally, it changes GTP binding activity, leading to autophosphorylation of LRRK2 protein, hyperphosphorylates kinases including MAPK, MMK2, MMK3, and MMK4^8^, and activates kinases like MMK1, with the ultimate consequence of cell death^9^. **Fig. 1** further illustrates a thorough explanation of the pathways impacted by the G2019S mutation^6,10–14^.Parkinsonism has also been linked to another significant mutation, L1120T, found in the kinase region of the LRRK2 protein, albeit the precise pathogenic mechanism causing it is not yet understood.

**Fig. 1:**
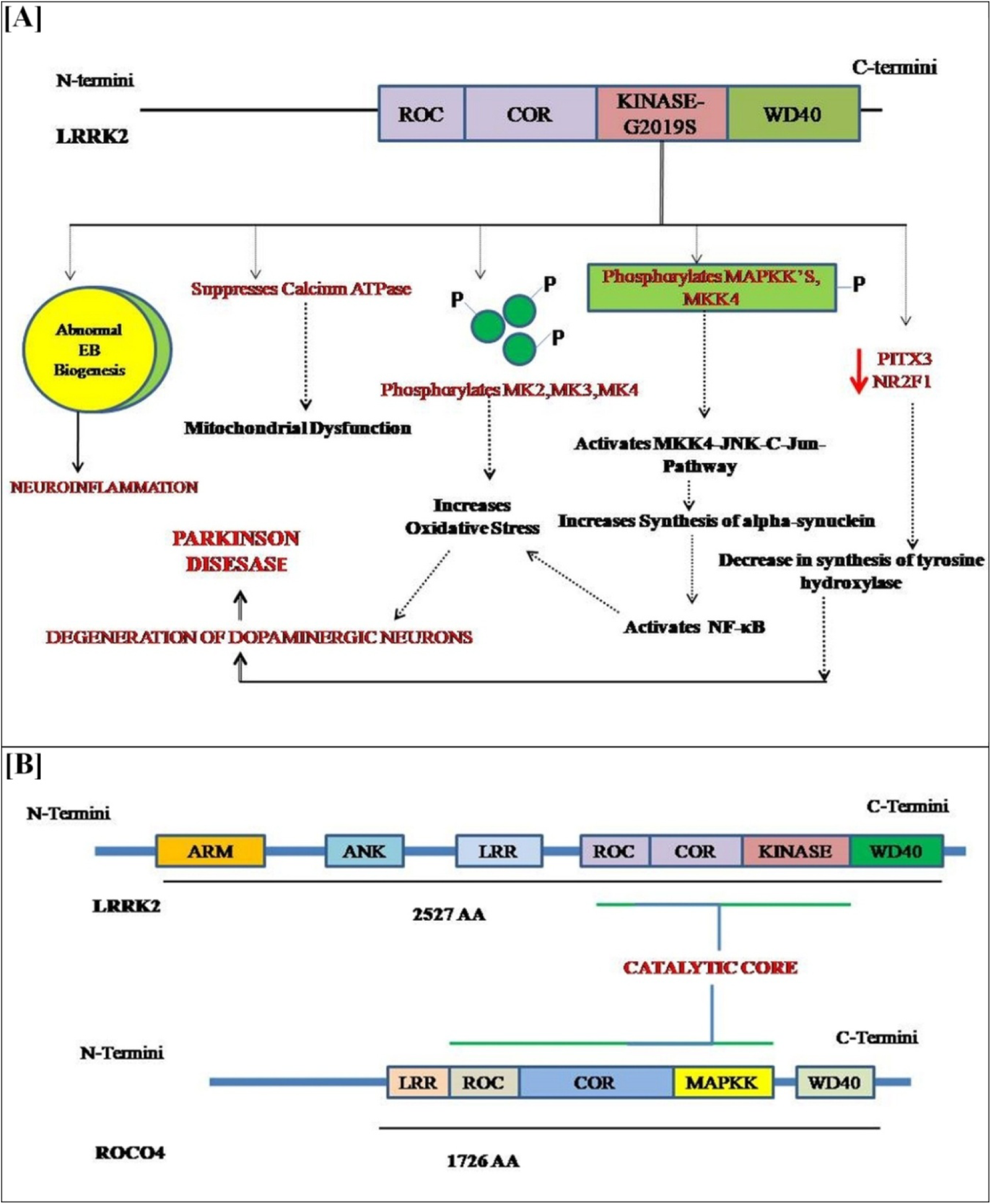
[a] Mechanism underlying G2019S mutation that causes Parkinsonism [b] Structure of LRRK2 and ROCO-4 kinase

The toxicity of G2019S mutation in LRRK2 is kinase-dependent, therefore designing or repurposing molecules that could inhibit the activity of kinase domain, possess the potential to confer neuro-protection and treat Parkinsonism. In clinical investigations employing rat models, synthetic compounds such as pyrrolo-pyrimidines and aryl benzamides have been used to suppress the kinase function of LRRK2 at the expense of lung and pulmonary toxicities^15^. However, using naturally existing plant-derived compounds is a substitute technique to replace synthetic medications and their accompanying toxicity. Phytochemicals from medicinal plants defend against oxidative damage, enhance cognitive health, maintain a proper balance of neurotransmitters, and are known to be less toxic^16^. Numerous phytochemicals, such as curcumin, resveratrol, berverin, terpenoids, and limonoids, have shown promise in the treatment of Parkinsonism and Alzheimer’s disease by binding to amyloid plaques or by regulating the expression of neurotrophic factors like nerve-growth factor (NGF), brain-derived neurotrophic factor (BNDF) and reducing neuroinflammation^17,18^.In light of this, phytochemicals represent a promising class of substances that may serve as treatments for a range of brain and neurodegenerative illnesses having multifactorial etiologies.

LRRK2 is a multi-domain Roco family-based protein in humans that exhibits similar characteristics and domain architecture to Roco4 protein present in *Dictyostelium discoideum*. In contrary to the difficulties faced to express and purify mammalian LRRK2 due to its enormous size of 2527 amino acid residues, a substantial amount of active and stable Roco4 protein could be easily expressed in E.coli. They are tractable, crystallizable, and the constructs for both wild and mutant variants in Roco 4 protein exhibit properties similar to those of mammalian LRKK2^19^.

To anticipate the similarity of the binding site between Roco 4 and LRRK2, mutant forms of Roco4 proteins were created by tweaking amino acid residues similar to the active site of human LRRK2 and referred to as “Humanized Roco4 kinase”^19^.Owing to similarities in its architecture and binding site with that of LRKK2, Humanized Roco 4 protein serves as an excellent structural framework to identify inhibitors to reverse the negative consequences of mutations linked to Parkinson’s disease.

Therefore, this study uses a curated library of ethano-botanical plant-derived chemicals for virtual screening and molecular dynamic simulation studies to uncover potential inhibitors of mutant forms of the kinase domain in LRRK2 protein, utilizing Humanized Roco4 as the model structure. This would further help us understand the potential of phytochemicals in treating Parkinson’s disease.

## Methodology

### Curation Of Phytochemicals

A literature review and database analysis was used to compile a list of 3069 phytochemicals identified in 100 plants. The scientific names of the plants used in this study are listed in **Table 1**. The plants were chosen in accordance with ethnobotanical records found in a variety of literatures, as well as with references found in texts on traditional medicine and literature reviews. The chemical structures for the phytochemicals were obtained from the open-source PubChem database (https://pubchem.ncbi.nlm.nih.gov/) and saved in .sdf format for further analysis.

**Table 1:**
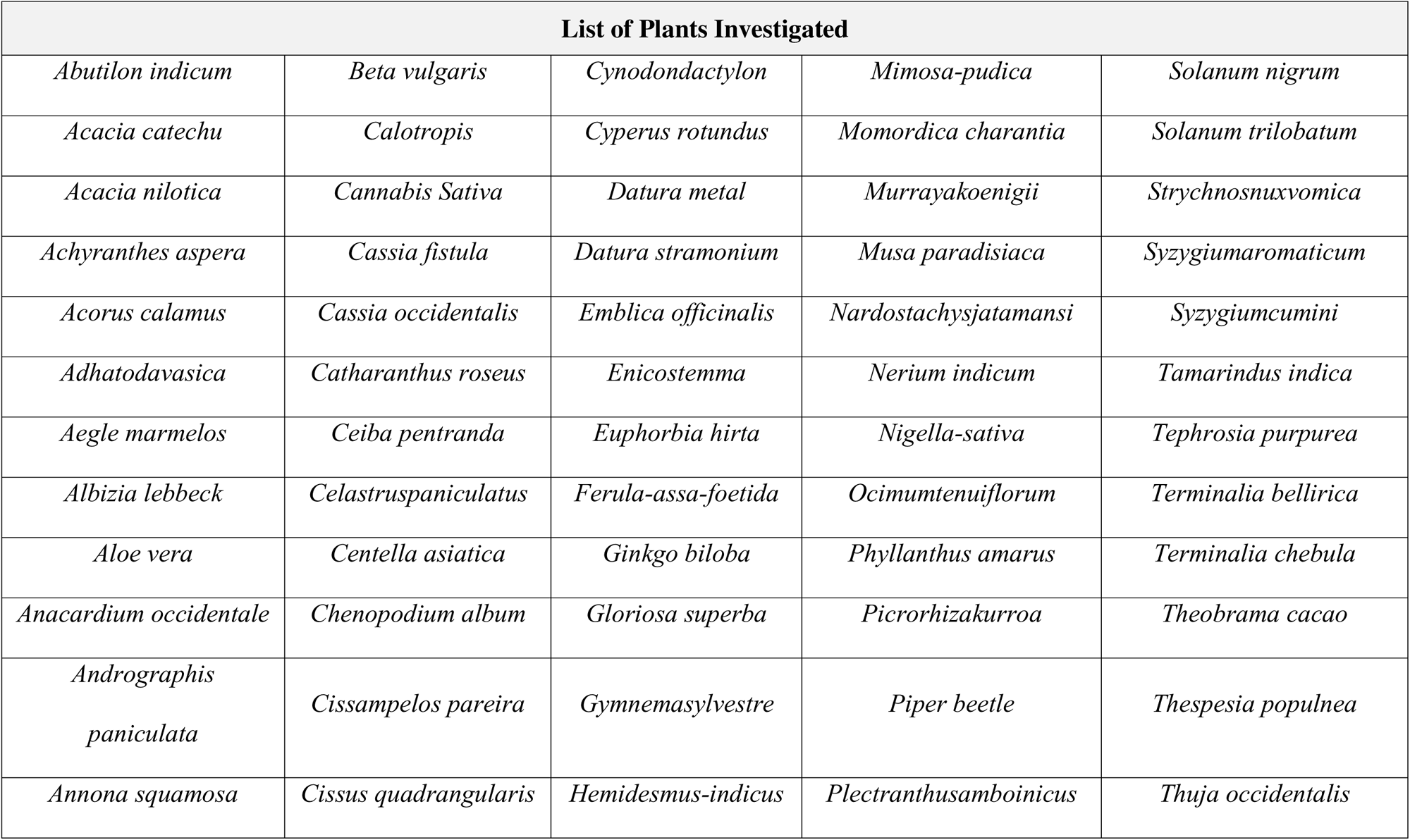
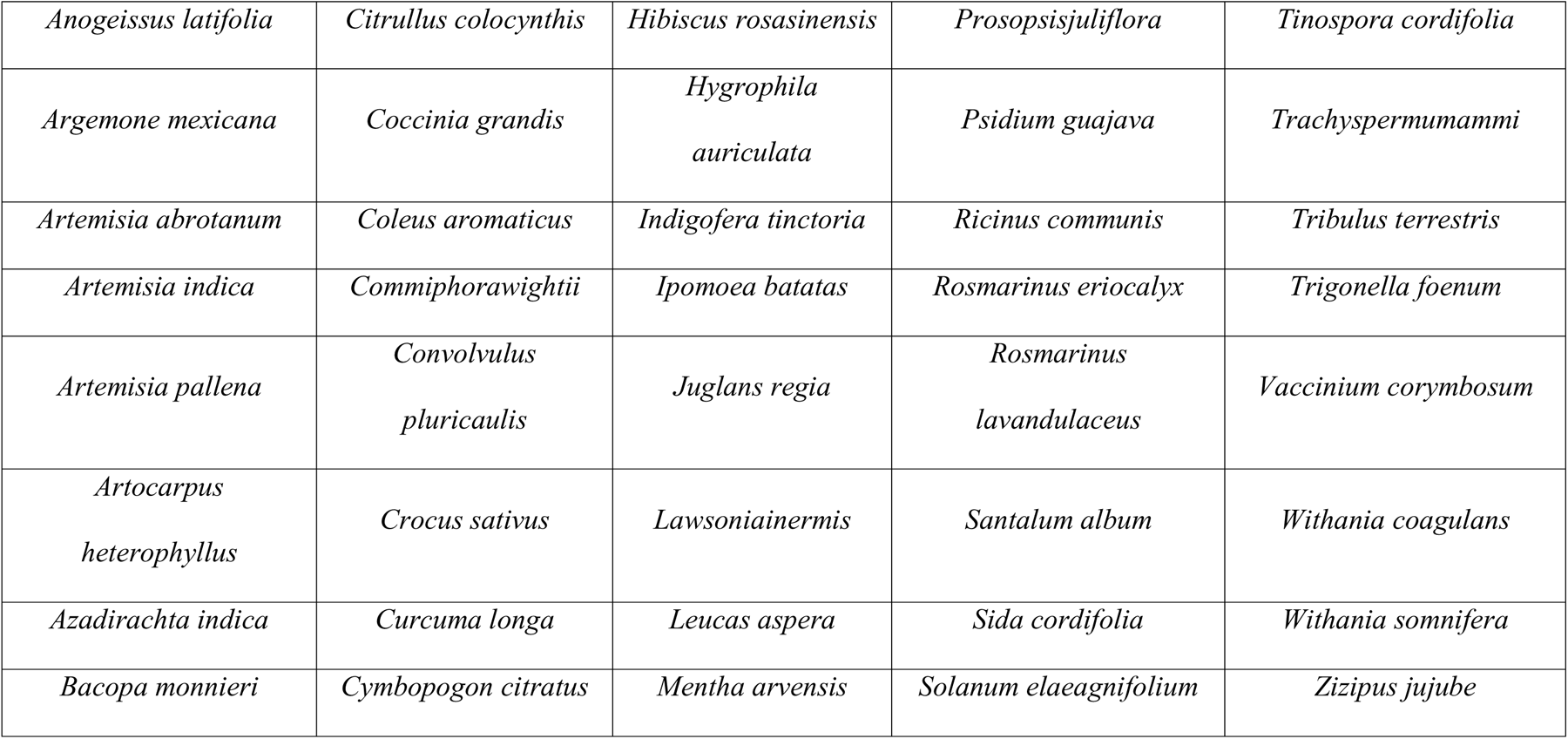
List of plants included in the current investigation.

| List of Plants Investigated |  |  |  |  |
| --- | --- | --- | --- | --- |
| *Abutilon indicum* | *Beta vulgaris* | *Cynodondactylon* | *Mimosa-pudica* | *Solanum nigrum* |
| *Acacia catechu* | *Calotropis* | *Cyperus rotundus* | *Momordica charantia* | *Solanum trilobatum* |
| *Acacia nilotica* | *Cannabis Sativa* | *Datura metal* | *Murrayakoenigii* | *Strychnosnuxvomica* |
| *Achyranthes aspera* | *Cassia fistula* | *Datura stramonium* | *Musa paradisiaca* | *Syzygiumaromaticum* |
| *Acorus calamus* | *Cassia occidentalis* | *Emblica officinalis* | *Nardostachysjatamansi* | *Syzygiumcumini* |
| *Adhatodavasica* | *Catharanthus roseus* | *Enicostemma* | *Nerium indicum* | *Tamarindus indica* |
| *Aegle marmelos* | *Ceiba pentranda* | *Euphorbia hirta* | *Nigella-sativa* | *Tephrosia purpurea* |
| *Albizia lebbeck* | *Celastruspaniculatus* | *Ferula-assa-foetida* | *Ocimumtenuiflorum* | *Terminalia bellirica* |
| *Aloe vera* | *Centella asiatica* | *Ginkgo biloba* | *Phyllanthus amarus* | *Terminalia chebula* |
| *Anacardium occidentale* | *Chenopodium album* | *Gloriosa superba* | *Picrorhizakurroa* | *Theobrama cacao* |
| *Andrographis*<br>*paniculata* | *Cissampelos pareira* | *Gymnemasylvestre* | *Piper beetle* | *Thespesia populnea* |
| *Annona squamosa* | *Cissus quadrangularis* | *Hemidesmus-indicus* | *Plectranthusamboinicus* | *Thuja occidentalis* |
| *Anogeissus latifolia* | *Citrullus colocynthis* | *Hibiscus rosasinensis* | *Prosopis juliflora* | *Tinospora cordifolia* |
| *Argemone mexicana* | *Coccinia grandis* | *Hygrophila*<br>*auriculata* | *Psidium guajava* | *Trachyspermum ammi* |
| *Artemisia abrotanum* | *Coleus aromaticus* | *Indigofera tinctoria* | *Ricinus communis* | *Tribulus terrestris* |
| *Artemisia indica* | *Commiphora wightii* | *Ipomoea batatas* | *Rosmarinus eriocalyx* | *Trigonella foenum* |
| *Artemisia pallens* | *Convolvulus*<br>*pluricaulis* | *Juglans regia* | *Rosmarinus*<br>*lavandulaceus* | *Vaccinium corymbosum* |
| *Artocarpus*<br>*heterophyllus* | *Crocus sativus* | *Lawsonia inermis* | *Santalum album* | *Withania coagulans* |
| *Azadirachta indica* | *Curcuma longa* | *Leucas aspera* | *Sida cordifolia* | *Withania somnifera* |
| *Bacopa monnieri* | *Cymbopogon citratus* | *Mentha arvensis* | *Solanum elaeagnifolium* | *Ziziphus jujube* |

### Drug Target

The kinase domain of ROCO4 protein, present in *Dictyostelium discoideum,* was chosen as the target protein due to its structural resemblance to LRRK2. The crystal structure and the FASTA sequence for the protein were obtained from Protein Data Bank (PDB ID: 4YZN). The downloaded protein structure was stripped of all non-amino acid residues (heteroatoms) and used further in the study. The 3D structures for the mutants G2019S and L1120T were generated using homology modeling via Swiss-Modeller (https://swissmodel.expasy.org/). The FASTA sequence of the protein was circumscribed by substituting the required alphabets (amino acid codes) at appropriate sites in the sequence to obtain the mutated primary structures. To obtain the 3D structures needed for virtual screening and MDS analysis, these structures were put through homology modeling. Three macromolecules were ultimately chosen for the analysis: the normal protein [Humanized ROCO 4], mutant-1 [Humanized ROCO 4 with the G2019S mutation], and mutant-2 [Humanized ROCO 4 with the L1120T mutation].

### Ligand and Protein Preparation

The Open babel tool in PyRx software was used to calculate the charge and minimize energy values for the ligands (phytochemicals). Energy minimization was performed with “uff force field” and “Conjugate Gradients” and the processed ligands were saved in pdbqt format for further analysis. The 3 protein drug targets (native protein, mutant 1, mutant 2) were prepared using the macromolecule preparation function in PyRx. The proteins were subjected to energy minimization and charge calculation, after which they were saved in .pdbqt format for further docking investigations.

### Identification of Active Site of Protein

PyMOL was used to visualize the three-dimensional structure of the ligand-protein interaction complex, PDB ID: 4YZN. The residues interacting with the co-crystallized ligand molecule (Compound 19), i.e., Gln-1031, Ile-1032, Gly-1033, Lys-1034, Val-1040, Lys-1042, ALA-1053, Lys-1055, Val-1091, Met-1105, Glu-1106, Leu-1107, Val-1108, Pro-1109, Cys-1110, Gly-1111, Asp-1112, Leu-1161, Ala-1176, and Asp-1177, were identified as the binding site/active site of the protein and were hence used for protein-ligand docking analyses in the virtual screening procedure.

### Virtual Drug-Screening of Phytochemical Ligands

Auto Dock Vina, present in PyRx, was used for virtual drug screening. The phytochemical ligand molecules were processed through ’Open Babel’ plug-in in PyRx. Energy minimization of ligands was performed using an “uff” force field and “conjugate gradient” algorithm. Target protein molecule “4YZN” was processed by removing all co-crystallized ligands, ions, and water molecules (heteroatoms) and was processed via the macromolecule preparation command in PyRx. The protein’s active site was covered by the grid box used for docking. Each of the target protein molecules was tested against all 3069 test ligands in a single run.

### Validation of Docking Protocol

The docking protocol and results were validated further by comparing the RMSD values of the co-crystal ligand and docked co-crystal ligand. The co-crystal ligand “Compound 19” in the target protein 4YZN was extracted and was subjected for protein-ligand docking via AutoDock Vina in PyRx tools. The co-crystal conformation and docked conformation were compared for their RMSD difference in the PyMOL tool for validation of the docking protocol. The docked conformation demonstrated less than 2.0Å RMSD with the co-crystallized conformation, hence confirming the reliability and validation of the docking protocol.

### Curation and Homology Modeling of ATP kinases

Using a literature search, a list of 34 kinases was compiled^20^.The Protein Data Bank was used to retrieve the crystal structures and FASTA sequences of the proteins. The downloaded protein structure was stripped off all non-amino acid residues and was further subjected to homology modeling wherever needed in order to obtain the 3D structures for virtual screening and determining the protein’s affinity toward the chosen ligands (Compound 19, Bacoside A, and Mongolicain A).

### Molecular-Dynamic Simulation

An intensive 100ns Molecular Dynamic Simulation (MDS) investigation was conducted using GROMACS running on the Ubuntu operating system (18.04 LTS). The phytochemicals Mongolicain A and Bacoside A that showed a substantial interaction with the Mutant-1 receptor molecules in docking study were then put through simulation experiments utilizing pyrrolopyrimidines (Compound 19 in 4YZN complex) as a reference ligand. To better understand the stability of the independent protein, the simulation was also run for each individual protein molecule. CHARMM-36 All Atoms force field, with TIP3P, was applied to carry out the procedures in simulation.

The receptors were organized into a GROMACS-recognizable structure file, and afterwards, topology was generated using the appropriate GROMACS commands. The CONF server was used to procure the structure file of ligands, and a Python script was used to create a GROMACS-recognizable structure file and construct the ligand topology. A protein-ligand structure file was then created by manually adding the parameters that were obtained from the construction of the ligand topology to the protein topology. The receptor-ligand complex was encased in a dodecahedron system of solvent molecules, and sodium ions were added in order to stabilize the system’s charge. Energy minimization, NVT, and NPT ensembles were executed according to the standard GROMACS simulation protocols provided in the webpage (http://www.mdtutorials.com/gmx/).

### ADMET Analysis of Ligands

The Adsorption, Distribution, Metabolism, Excretion, and Toxicity (ADMET) and Drugability for the test and reference ligands were investigated using Molinspiration (https://molinspiration.com/) andpreADMET (https://preadmet.qsarhub.com)

## Results

### Target Protein & Binding Site

The drug target focused in the study was the kinase domain of the ROCO 4 protein. The crystal structure of Humanized ROCO4 protein was retrieved from the PDB database (PDB ID: 4YZN) and was used as a native protein. The primary structure of the protein was altered and homology modeling was applied to generate the necessary mutant variants, G2019S and L1120T, linked to Parkinsonism in humans. The homology model of the G2019S mutant is shown in **Fig. 2**, which highlights the binding site and mutation site along with a Ramachandran plot that reveals 96.32 percent of the residues are present in the favorable region. The homology model of the L1120T mutant also demonstrated a similar Ramachandran analysis, with 96.32 percent of residues being present in the favorable region.

**Fig. 2:**
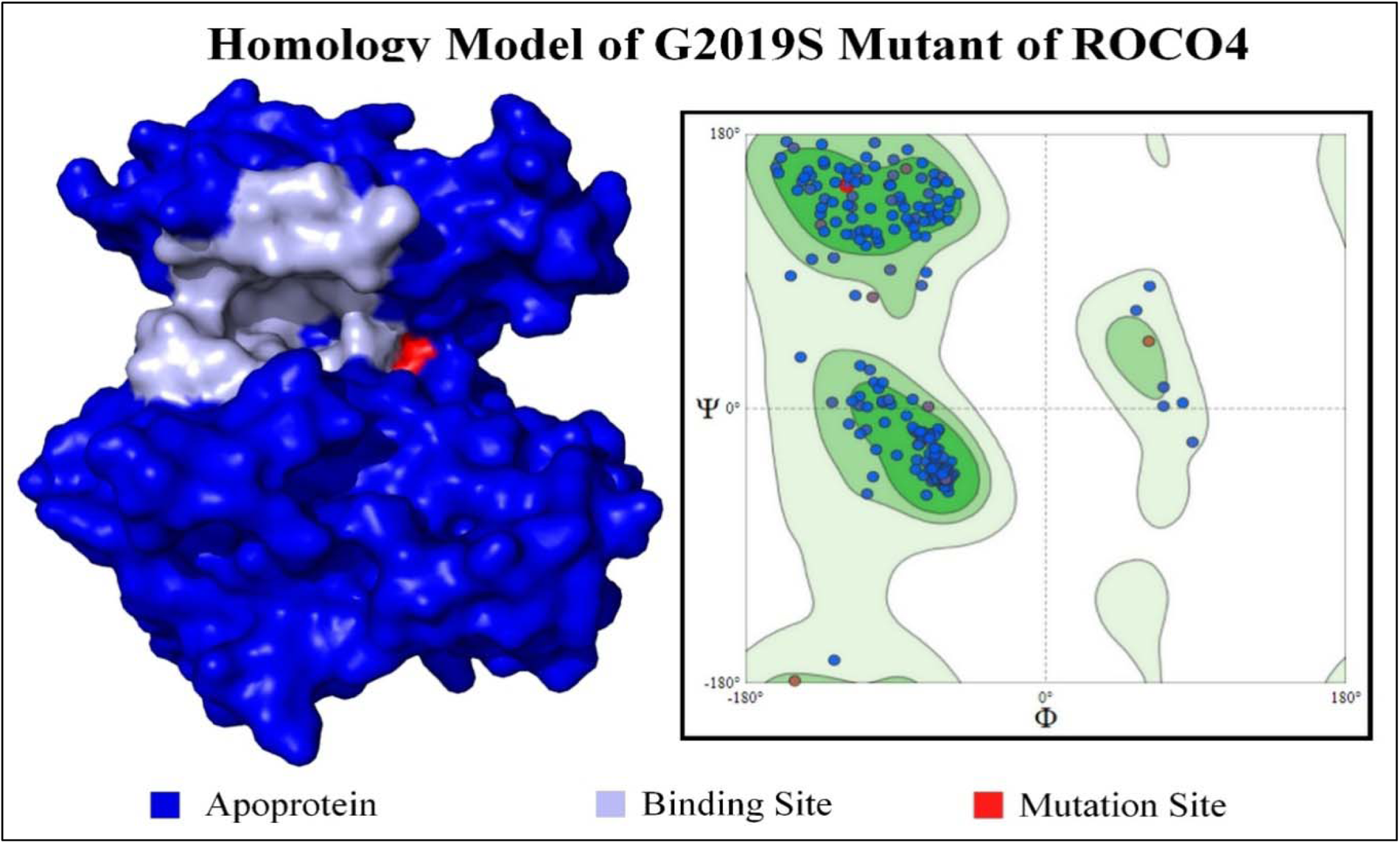
Homology Model of G2019S Mutant (Mutant-1) of ROCO4 with labeled binding site, mutation site and the Ramachandran Plot

Based on the interactions of the co-crystallized ligand Compound 19 in 4YZN structure, the binding site of the protein was identified as Gln-1031, Ile-1032, Gly-1033, Lys-1034, Val-1040,Lys-1042, Ala-1053, Lys-1055,Val-1091,Met-1105, Glu-1106, Leu-1107, Val-1108, Pro-1109, Cys-1110, Gly-1111, Asp-1112, Leu-1161, Ala-1176, andAsp-1177. The grid box was placed in the identified binding locations for all 3 proteins to conduct a protein-ligand docking investigation.

### Protein-Ligand Docking with Target Protein

The Virtual Drug Screening protocol was performed using Auto Dock Vina, in-built within the PyRx software. A total of 3069 phytochemicals were curated from 100 recognized plant sources. A final set of 1795 phytochemicals were subjected to protein-ligand docking after recurring ligands and molecules those with a molecular weight of less than 200 g/mol were removed. The protein-ligand docking was performed against 3 macromolecules, i.e., native protein, mutant-1, and mutant-2. The docking study was duplicated (re-docking) to check for reproducibility and accuracy of the binding energy. The results of the protein-ligand docking analyses are tabulated and compared in **Table 2**, which shows the top scoring phytochemical hits for all three macromolecules.

**Table 2:** Results of Protein-Ligand docking analysis for top 10 Ligand Hits.

| PubChem | Phytochemical Name | Wild Type Protein |  | Mutant 1 |  | Mutant 2 |  |
| --- | --- | --- | --- | --- | --- | --- | --- |
|  |  | Docking<br>(Kcal/mol) | Redocking<br>(Kcal/mol) | Docking<br>(Kcal/mol) | Redocking<br>(Kcal/mol) | Docking<br>(Kcal/mol) | Redocking<br>(Kcal/mol) |
| 92043183 | Bacoside-A | -11.3 | -11.3 | -11.2 | -11.4 | -11 | -11.3 |
| 78172774 | Guavin B | -10.7 | -10.8 | -10.8 | -10.8 | -10.6 | -10.6 |
| 16099532 | Ashwagandha Bolide | -10.5 | -10.5 | -10.5 | -8.6 | -9.6 | -9.5 |
| 442985 | Solasodine | -10.2 | -10.2 | -10.3 | -10.3 | -9 | -9 |
| 91472 | Friedelin | -10.2 | -10.2 | -10.2 | -10.2 | -9.4 | -9.3 |
| 99474 | Diosgenin | -10.2 | -10.2 | -10.2 | -10.2 | -8.9 | -8.9 |
| 73814547 | Guavin D | -10.1 | -10.1 | -10.2 | -10.2 | -10.6 | -10.7 |
| 12304430 | Episarsasapogenin | -10.1 | -10.1 | -9.9 | -10.1 | -9.3 | -9.4 |
| 16131089 | Vescalagin Carboxylic Acid | -10 | -10 | -10 | -10 | -10.2 | -10.2 |
| 6737485 | Bianthraquinone | -9.9 | -9.8 | -9.9 | -9.9 | -10.5 | -10.5 |
| 16167195 | Acutissimin A | -9.8 | -9.8 | -9.8 | -9.8 | -10.8 | -10.8 |
| 11766372 | Tellimagrandin II | -9.8 | -9.8 | -9.8 | -9.8 | -10.3 | -10.3 |
| 101618906 | Gypenoside XXVIII | -9.7 | -9.8 | -9.8 | -9.7 | -10.6 | -10.6 |
| 156784 | GRECUNIN G | -9.6 | -9.7 | -9.6 | -9.4 | -10.4 | -10.4 |
| **44576149** | **Mongolicain-A** | **-9.3** | **-9.3** | **-12.3** | **-12.3** | **-8.3** | **-8.3** |
| 101676952 | Eugenigrandin A | -9.2 | -10 | -10.1 | -10.1 | -10.5 | -10.5 |
| 85348461 | Centella Saponin C | -8.9 | -8.9 | -8.9 | -8.9 | -10.4 | -10 |
| Compound 19 | Pyrolopyrimidine | -7.7 | -7.7 | -7.8 | -7.8 | -7.6 | -7.6 |

Among the listed phytochemicals, the most significant interaction was demonstrated by Mongolicain-A having a binding energy of -12.3 kcal/mol, followed by the formation of 8 hydrogen bonds (Lys-29, Lys-37, Lys-50, Gly-106, and Asp-172) with G2019S Mutant (Mutant-1). Mongolicain-A demonstrated a very high specificity towards Mutant-1 in comparison to the native and Mutant-2 forms of the protein. This specificity was observed in re-docking as well, confirming the high selective affinity of Mongolicain-A towards Mutant-1 of the ROCO4 protein. Bacoside-A had the highest affinity after Mongolicain-A, with a binding energy of -11.4 kcal/mol, followed by the formation of 6 hydrogen bonds with the mutant 1 protein. Bacoside-A, on the other hand, exhibited no specificity or selectivity to either form of ROCO4 protein, with a binding energy of -11.0 or greater for all three macromolecules studied. The reference molecule, i.e., Compound 19 (Pyrrolopyrimidines), a co-crystallized inhibitor of Humanized ROCO4 protein (PDB ID: 4YZN), demonstrated a much lower affinity towards the target molecules, with a binding energy of -7.8 kcal/mol following the formation of 3 hydrogen bonds with Gly-31 and Asp-107. This was much below the range of binding energy exhibited by the phytochemicals Mongolicain-A and Bacoside-A. Graphical representations of the 3 protein-ligand docking complexes, i.e., Mutant-1 with Compound-19, Bacoside-A, and Mongolicain-A, are represented in **Fig. 3**.

**Fig. 3:**
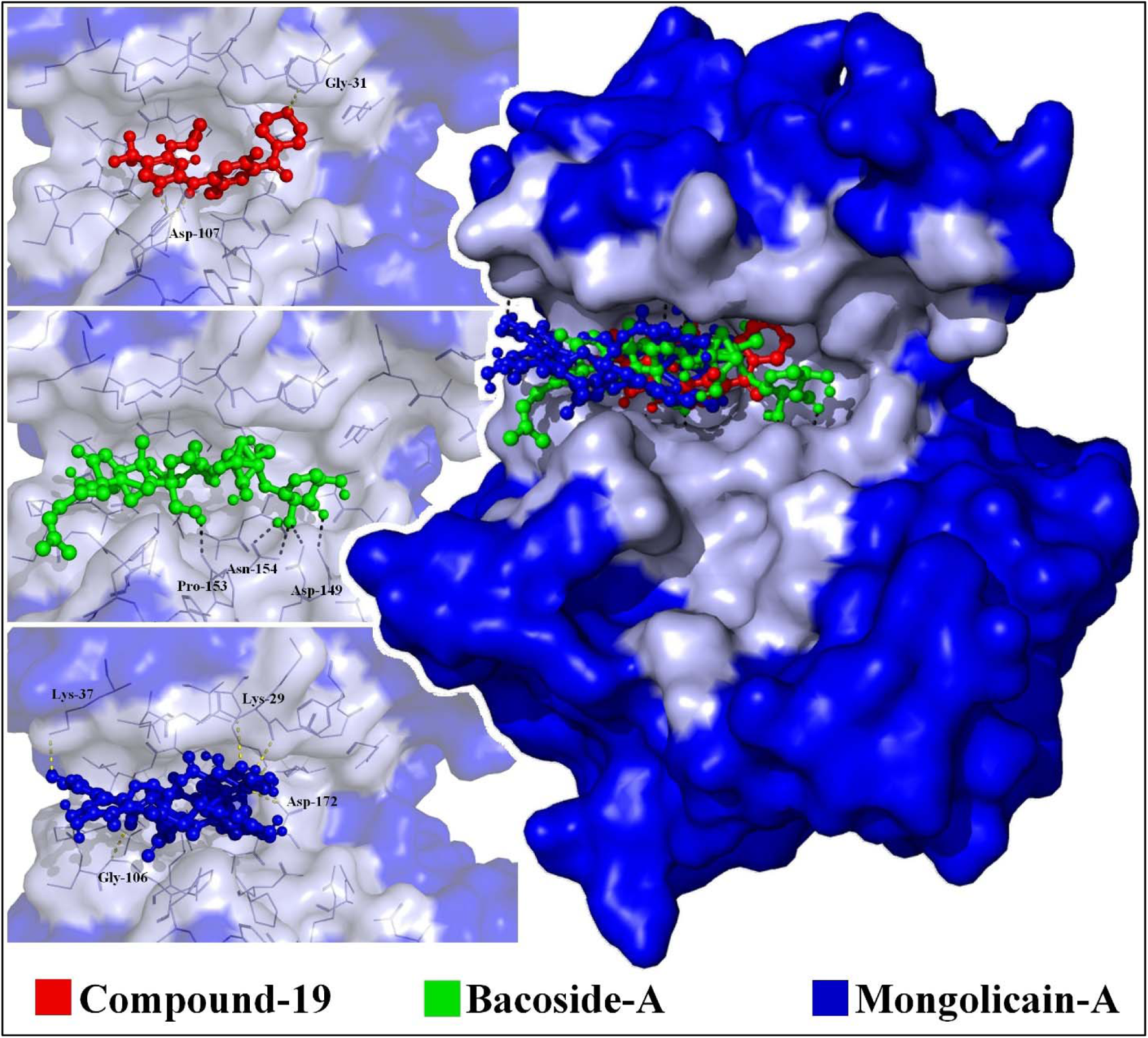
Interaction of Mutant-1 with Compound-19, Bacoside-A and Mongolicain-A

Mongolicain-A and Bacoside-A, the two identified phytochemicals, are seen to be enriched with hydroxyl groups, keto groups, and aromatic rings. These functional groups present in the molecules provide a major contribution to their polar interaction potential, which has a considerable impact on the complex’s binding potential and protein-ligand interaction stability. Hence, based on protein-ligand docking analyses, the two phytochemical ligands (Mongolicain-A and Bacoside-A) were identified as effective kinase inhibitors of the G2019S mutant (Mutant-1). These molecules were further subjected to Molecular Dynamics Simulation (MDS) studies.

### Protein-Ligand Docking with Kinases

A protein-ligand docking analysis was performed on 34 different kinase proteins to test the specificity and affinity of the ligands, namely Bacoside A, Mongolicain A, and Compound 19, against other Kinase family proteins. The results of the kinase specificity docking analysis are tabulated in **Table 3**.

**Table 3:**
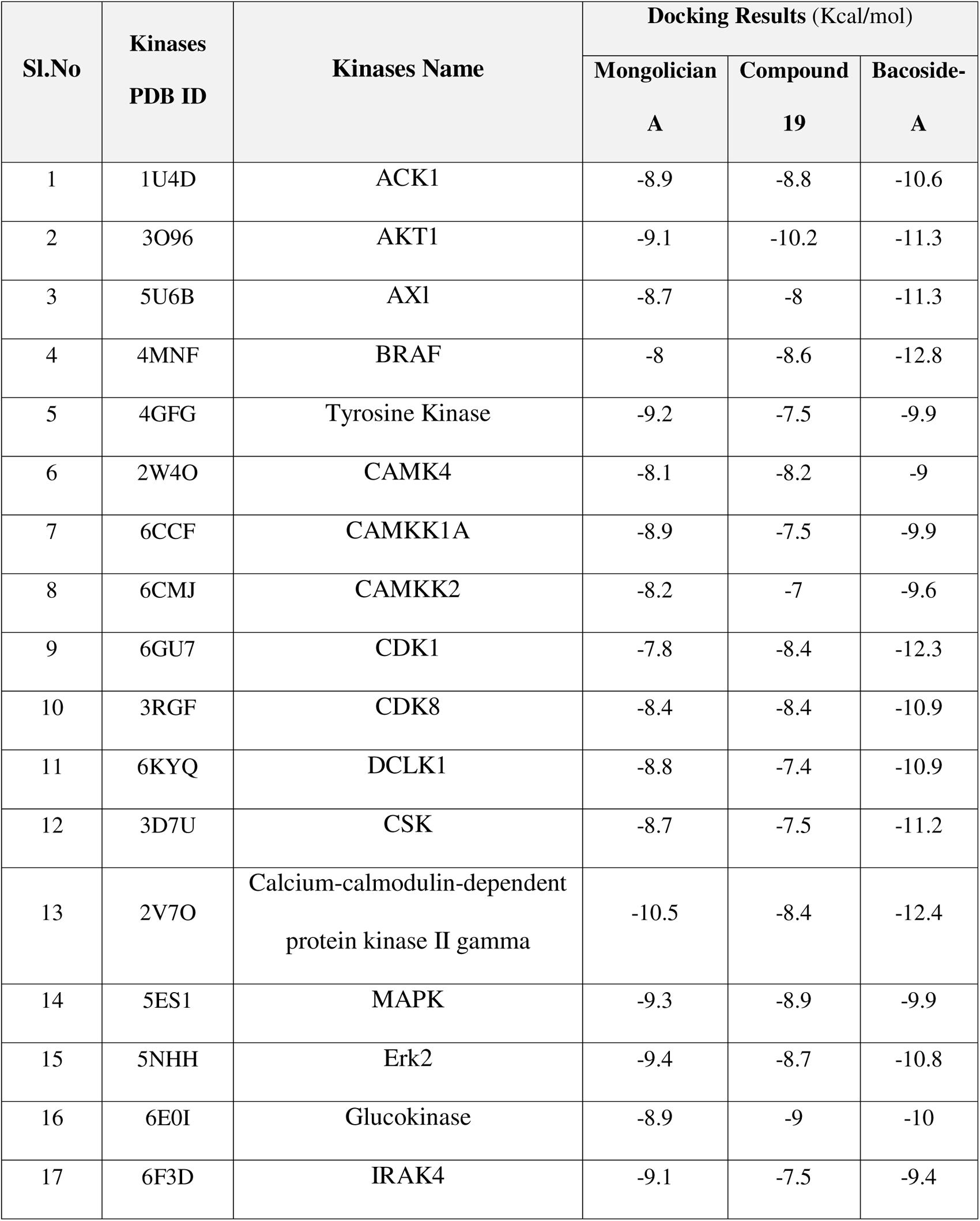
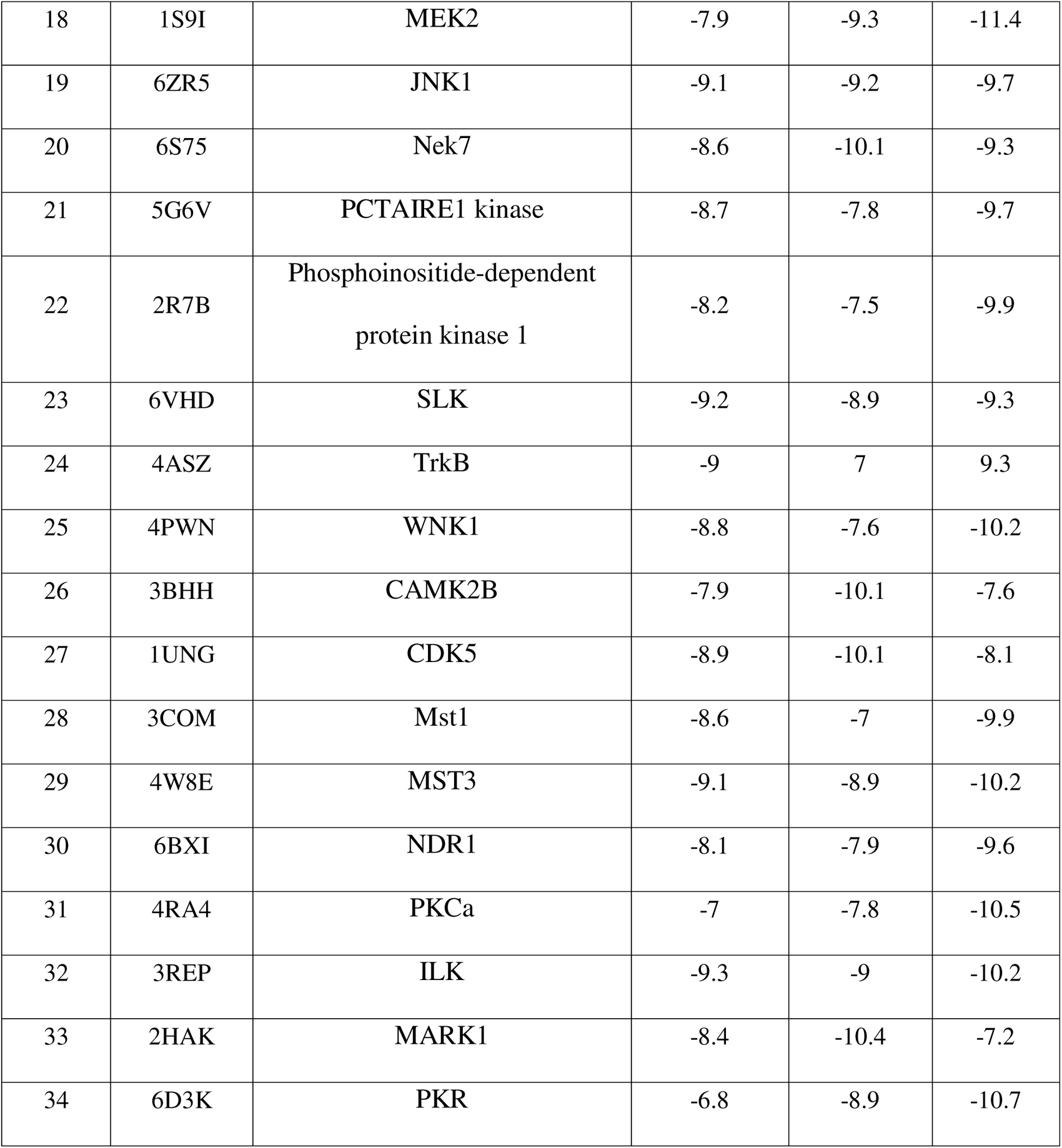
Results of Kinase Specificity Protein-Ligand Docking analysis with Other Family Kinases (Human)

Among the 34 kinase proteins, the test ligand Mongolicain-A demonstrated a highest affinity of -10.5 kcal/mol against the “Calcium-calmodulin-dependent protein kinase II gamma” protein (PDB ID: 2V7O), which was much lower than the affinity towards the Mutant-1 protein (-12.3 kcal/mol). Due to this reduced affinity towards other kinase family proteins and also to the native form of the ROCO4 protein itself, it was further justified that Mongolicain-A is highly specific to Mutant-1 of the ROCO4 protein and that it would exhibit low or no side-effects due to its lesser affinity to other similar kinase proteins.

Bacoside-A demonstrated a significant binding affinity to all the kinases, with the highest affinity of -12.8 kcal/mol against “CDK1” protein (PDB ID: 6GU7). This further confirms that Bacoside-A is an effective kinase inhibitor, but it lacks specificity to any individual protein but rather holds potential to function as a general kinase inhibitor. Surprisingly, the reference ligand Compound-19 demonstrated significant binding potential with all the tested kinase proteins, and the highest affinity of -10.4 kcal/mol was observed against the “MARK1” protein (PDB ID: 2HAK). This affinity was much higher than the affinity demonstrated against the target ROCO4 proteins.

Overall, Mongolicain-A showed good potential as a selective ROCO4 kinase mutant-1 inhibitor. Quantitative affinity values strongly indicate that Mongolicain A had very high affinity towards the targeted ROCO4 mutant-1 protein. Despite the considerable affinity of Mongolicain-A towards other kinases, it demonstrated an exceptional affinity to the Mutant-1 of ROCO-4 protein, suggesting it to be a selective and specific inhibitor of the target. Whereas, Bacoside-A was identified as broad-spectrum kinase inhibitor, since Bacoside-A demonstrates significant inhibition potential with all the kinase targets investigated in this study.

### Molecular Dynamics Simulation Analysis

To investigate the protein-ligand complex stability and inhibition potential of test ligands “Mongolicain-A” and “Bacoside-A” against the target protein “Mutant-1 of ROCO4”, an exhaustive Molecular Dynamics Simulation (MDS) of 100 ns was performed using GROMACS. Four individual simulations were executed, i.e., (i) Mutant-1 Protein individually; (ii) Mutant-1 in complex with Compound-19; (iii) Mutant-1 in complex with Bacoside-A; (iv) Mutant-1 in complex with Mongolicain-A.

The results of the MDS analyses are represented and discussed in terms of their RMSD, RMSF, SASA, Potential energy, and Hydrogen bond plots.

### Root Mean Square Deviation (RMSD) and Root Mean Square Fluctuation (RMSF)

The stability of the protein-ligand complex and protein structures are measured in terms of the RMSD plot. The comparison of RMSD values of all 4 MDS are represented in Fig. 4a. All 4 MDS exhibited significant stability over the 100 ns period of simulation, with an average of 0.15 nm RMSD. Throughout the simulation period, all four complexes were found to be very stable, with no significant fluctuations in their RMSD..Investigation of the highest RMSD value for individual MDS complexes suggested that Bacoside-A demonstrated 0.267 nm, Mongolicain-A demonstrated 0.222 nm, and Compound-19 demonstrated 0.238 nm RMSD. Based on the RMSD plot analyses, it was evident that all 4 MDS complexes were very stable, and all 3 ligands, i.e., Mongolicain-A, Bacoside-A, and Compound-19, demonstrate strong potential as inhibitors of Mutant-1 of the ROCO4 protein.

**Fig. 4:**
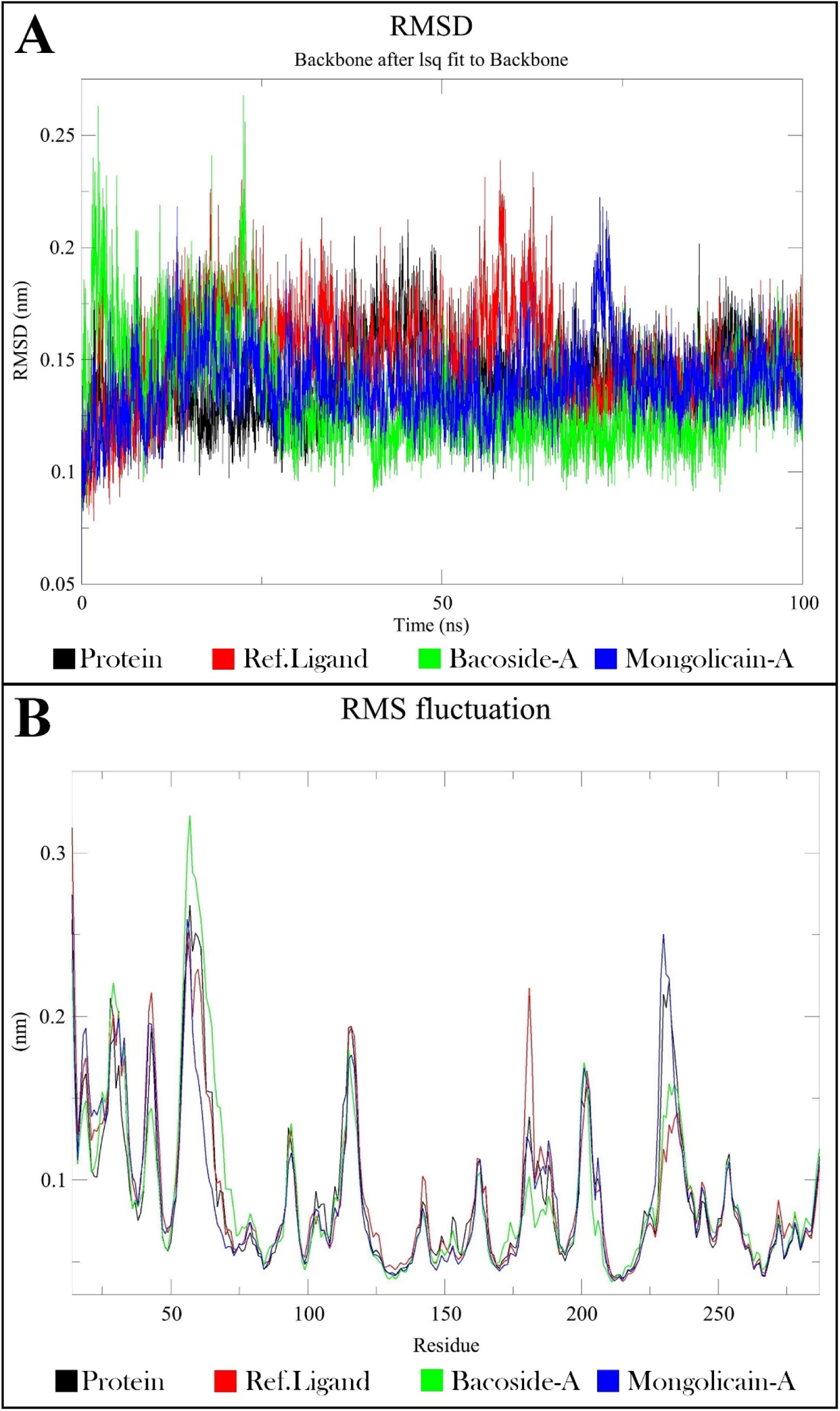
Molecular Dynamic Simulation Results; a) RMSD plot; b) RMSF plot

RMSF plots of individual amino acid residues in all 4 MDS complexes are compared in Fig. 4b. The majority of the amino acid residues demonstrated less than 0.1 nm fluctuation, with the exception of a few residues with fluctuations above 0.2 nm. Analyses of individual MDS complexes show that, highest fluctuation was demonstrated by Ser-57 (0.32 nm) in the Bacoside-A complex, by Pro-14 (0.274 nm) in the Mongolicain-A complex, and by Pro-14 (0.315 nm) in the Compound-19 complex. Interestingly two amino acid residues, Ser-182 and Ser-184, that play an important role in maintaining the kinase activity of ROCO4 kinase through autophosphorylation *(Kortholt et al., 2012),* had demonstrated significantly lower RMSF values for test ligands. Ser-182 and Ser-184 demonstrated a RMSF value of 1.143Å and 0.995Å for Mongolicain A, and 0.858Å and 0.743Å for Bacoside A. Whereas Ser-182 and Ser-184 demonstrated RMSF values of 1.716Å and 1.058Å for Compound 19 respectively. The RMSF analyses, however, strongly suggest that the four MDS complexes are stable and that the two test ligands are potential inhibitors of the chosen target protein.

### Solvent Accessible Surface Area (SASA)

SASA analysis can be used to comprehend the stability of the protein-ligand complex and the inhibition potential of the ligand. Fig. 5a compares the SASA assessments of four MDS complexes. Overall observation revealed that Mongolicain-A exhibits an average SASA of 150 nm2, while Bacoside-A exhibits the lowest SASA throughout the simulation with an average value of 145 nm2. Bacoside-A was observed to have much higher stability with the target protein than Mongolicain-A and Compound-19. Overall findings, however, revealed that all 3 protein-ligand complexes displayed significant stability throughout the 100 ns simulation.

**Fig. 5:**
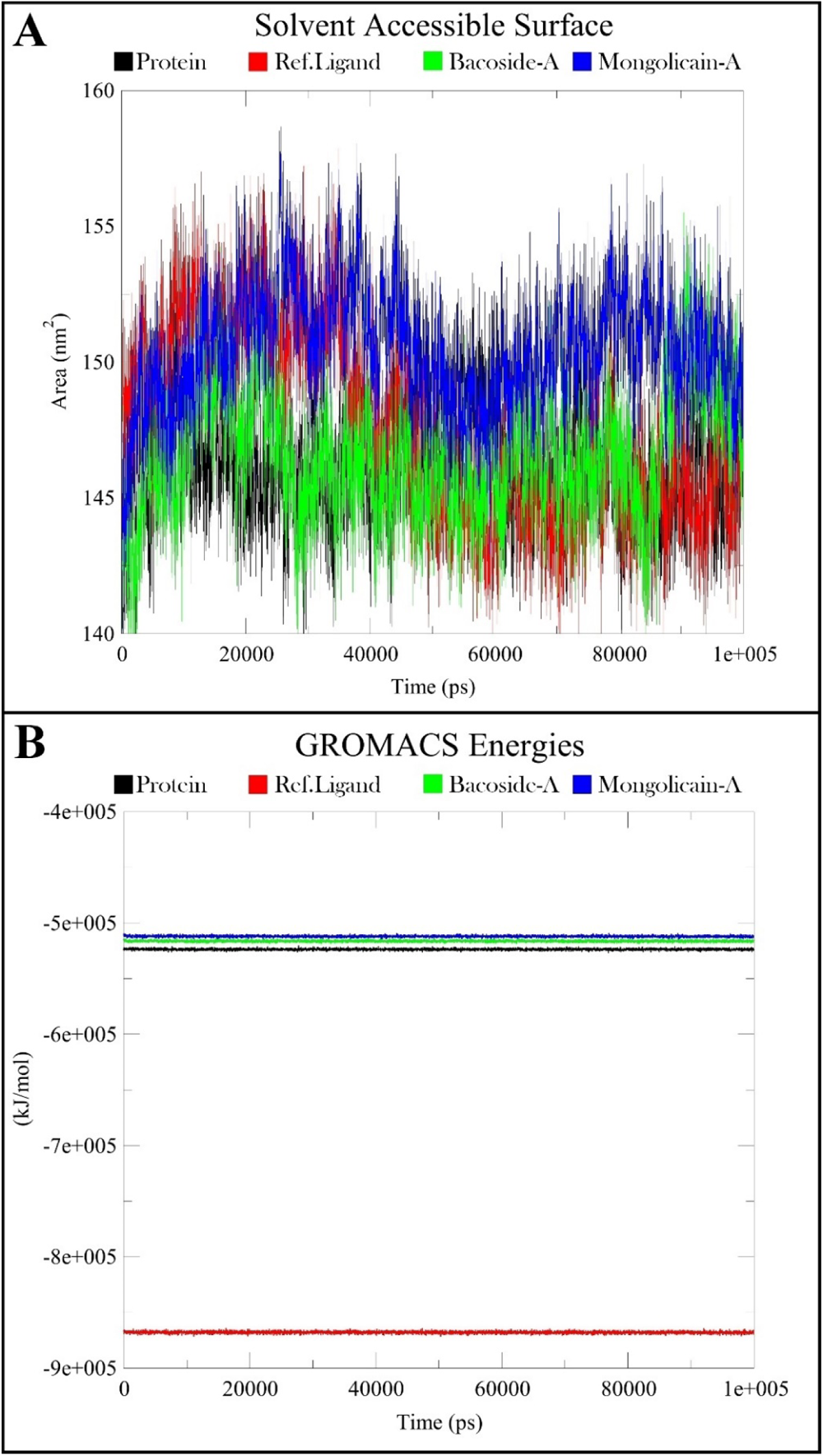
Molecular Dynamic Simulation Results; a) SASA plot; b) Potential Energy plot

### Potential Energy

Potential energy plots of all 4 MDS are shown in Fig. 5b. The potential energy of the target protein was significantly reduced when it was complexed with Mongolicain-A and Bacoside-A. However, Compound-19 significantly increased the potential energy of the complex. Mutant-1 of the ROCO4 protein demonstrated the lowest potential energy of -526955.93 kJ/mol at 76.93 ns. This energy was further reduced when complexed with the test ligands, i.e., Mongolicain-A demonstrated a lowest potential energy of -515056.71 kJ/mol at 87.99 ns, and Bacoside-A demonstrated a lowest potential energy of -519015.28 kJ/mol at 41.88 ns. Compound-19 increased the potential energy of the complex and demonstrated the lowest potential energy of -871675.18 kJ/mol at 50.27 ns. The protein ligand complex for both test ligands, Mongolicain-A and Bacoside-A, was determined to be stable and capable of acting as a target protein inhibitor based on potential energy studies.

### Hydrogen Bond & Interaction Analysis

Stability and strength of the protein-ligand complex were also analyzed in terms of the number of hydrogen bonds exhibited during the simulation period and the number of close-range proximity residues. **Fig. 6** depicts a graphical representation of the number of hydrogen bonds and close-range proximities interacting residues for all three protein-ligand complexes. Initial observation clearly indicates that the maximum numbers of interactions are formed by Mongolicain-A, with an average of at least 4 hydrogen bonds (over 75% of frames) and a maximum of 8 hydrogen bonds with the protein. Next to this, Bacoside-A formed an average of 2 hydrogen bonds (over 50% frames) and a maximum of 3 hydrogen bonds. The lowest number of hydrogen bonds was formed by Compound-19 with an average of 1 hydrogen bond (over 50% frame) and a maximum of 3 hydrogen bonds. The total number of hydrogen bonds indicates the strength of the polar interactions, and the number of proximity residues (less than 0.35 nm distance) denotes the strength of the hydrophobic or non-polar interactions. Based on the interaction analysis of the 3 complexes, it was confirmed that, among the 3 complexes, Mongolicain-A demonstrates the highest potential as an inhibitor.

**Fig. 6:**
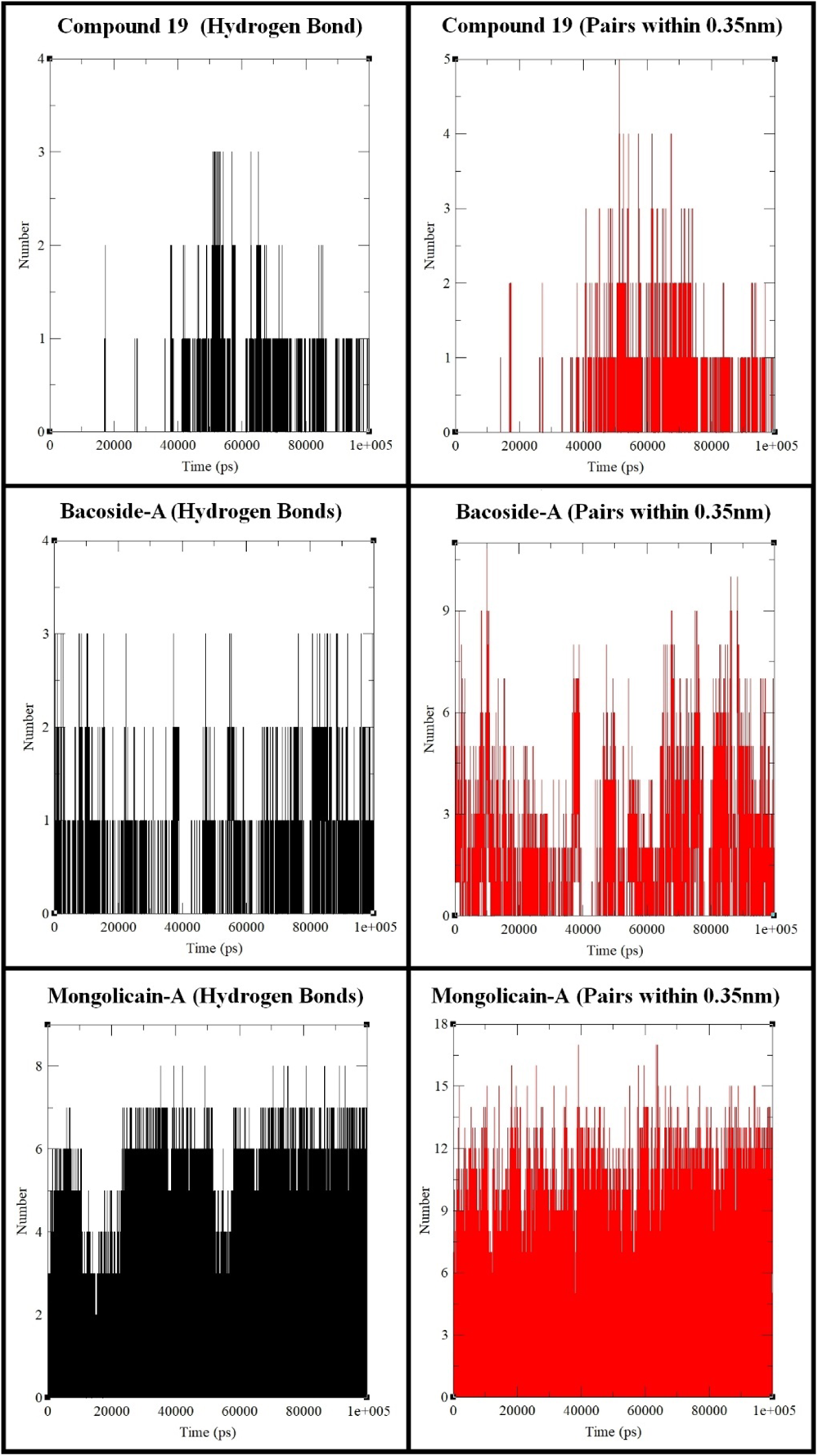
Molecular Dynamic Simulation Results as Hydrogen Bond Analysis

### ADMET & Drugability Analysis

The pre-ADMET and Molinspiration webtools were used to analyze the absorption, distribution, metabolism, excretion, and toxicity of the two test phytochemical ligands (Mongolicain A and Bacoside A) and the reference molecule (Compound 19). Results of the in-silico ADMET and Drugability analyses of the three ligand molecules are summarized in Table 4. All the physicochemical properties of the three ligands were favorable as a drug-like molecule.

**Table 4:**
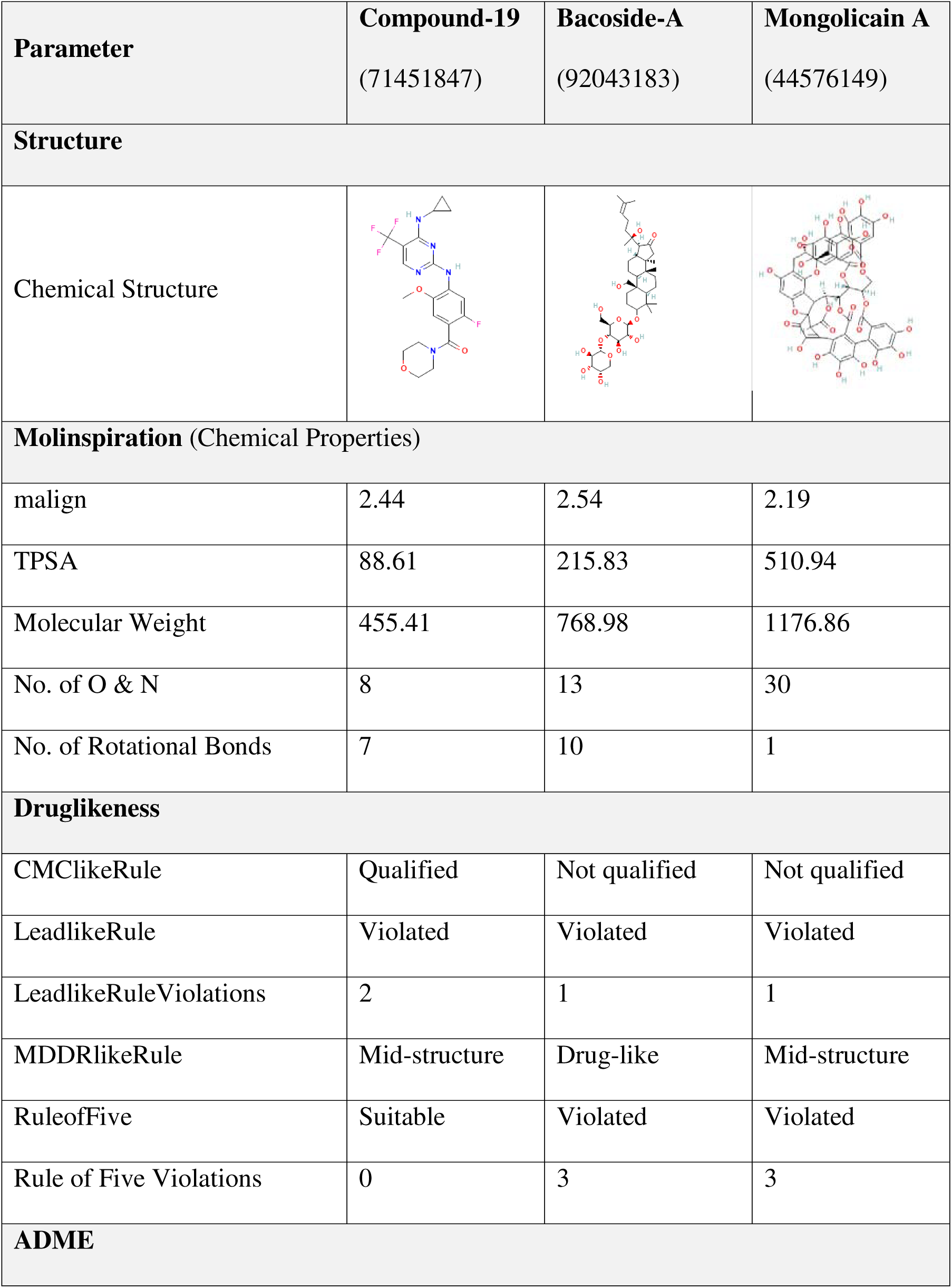
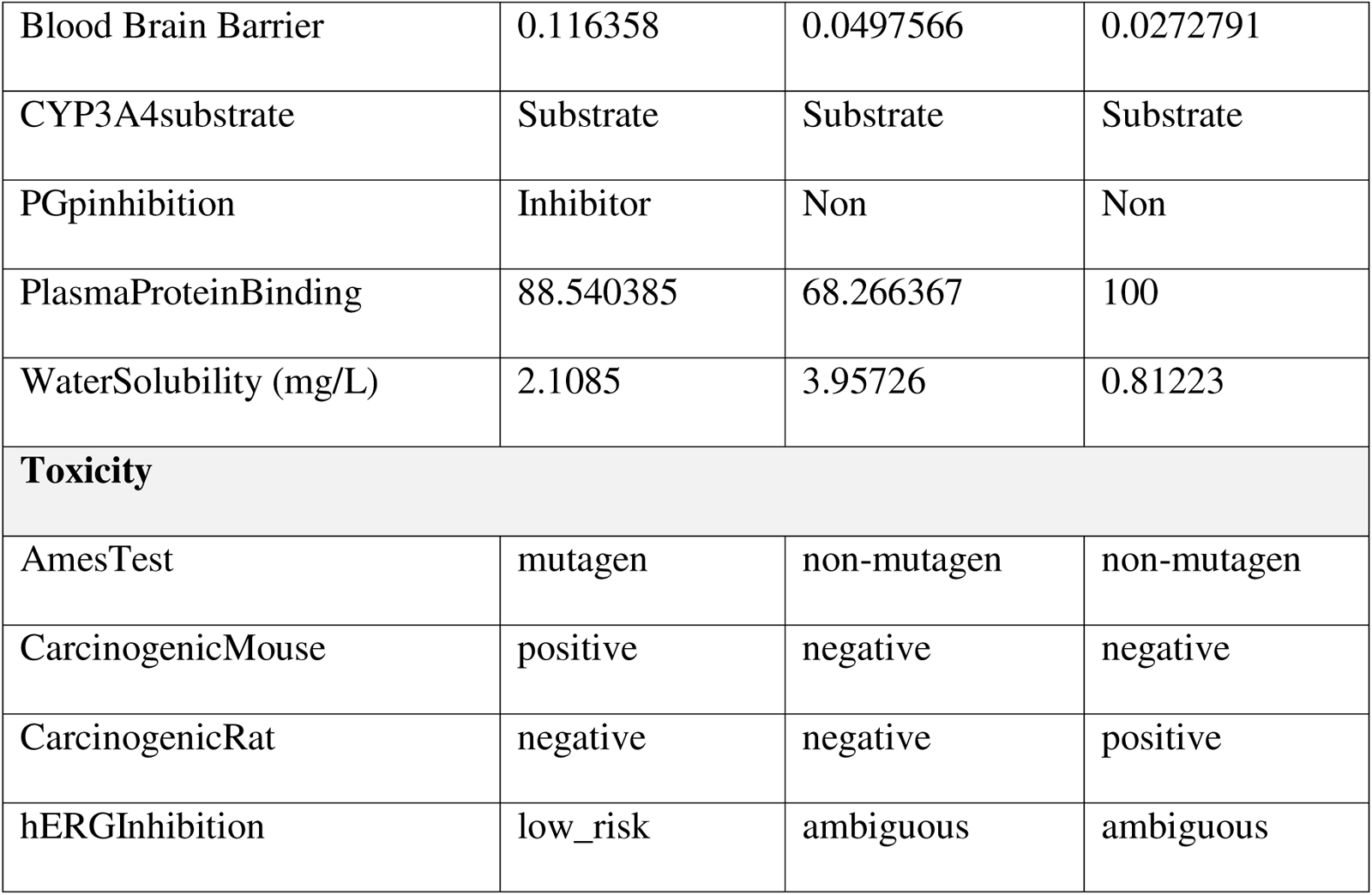
ADMET / Drugability analysis of target ligand molecules.

| Parameter | Compound-19<br>(71451847) | Bacoside-A<br>(92043183) | Mongolicain A<br>(44576149) |
| --- | --- | --- | --- |
| **Structure** |  |  |  |
| Chemical Structure |  |  |  |
| **Molinspiration (Chemical Properties)** |  |  |  |
| malign | 2.44 | 2.54 | 2.19 |
| TPSA | 88.61 | 215.83 | 510.94 |
| Molecular Weight | 455.41 | 768.98 | 1176.86 |
| No. of O & N | 8 | 13 | 30 |
| No. of Rotational Bonds | 7 | 10 | 1 |
| **Druglikeness** |  |  |  |
| CMClkeRule | Qualified | Not qualified | Not qualified |
| LeadlikeRule | Violated | Violated | Violated |
| LeadlikeRuleViolations | 2 | 1 | 1 |
| MDDRlikeRule | Mid-structure | Drug-like | Mid-structure |
| RuleofFive | Suitable | Violated | Violated |
| Rule of Five Violations | 0 | 3 | 3 |
| **ADME** |  |  |  |
| Blood Brain Barrier | 0.116358 | 0.0497566 | 0.0272791 |
| CYP3A4substrate | Substrate | Substrate | Substrate |
| PGpinhibition | Inhibitor | Non | Non |
| PlasmaProteinBinding | 88.540385 | 68.266367 | 100 |
| WaterSolubility (mg/L) | 2.1085 | 3.95726 | 0.81223 |
| **Toxicity** |  |  |  |
| AmesTest | mutagen | non-mutagen | non-mutagen |
| CarcinogenicMouse | positive | negative | negative |
| CarcinogenicRat | negative | negative | positive |
| hERGIInhibition | low_risk | ambiguous | ambiguous |

The two test ligands do violate the few drug likeness rules, but as the molecules are natural, plant-derived products, these violations are still acceptable. All three molecules have low blood-brain barrier permeability and are easily metabolized and excreted from cells without causing major side effects. Toxicity predictions of the ligands suggest that the test ligands show the least side-effects, with their mutagenic and carcinogenic properties being present in an acceptable range.

In-silico ADMET and Drug likeness analysis of the ligands suggests that these molecules are suitable drug-like candidates and can be further probed to be applied for human applications in treating Parkinsonism.

## Discussion & Conclusion

According to the results of the Molecular Dynamic Simulation study, Mongolicain-A, which is derived from *Psidium guajava* (Guava), may be a possible selective inhibitor against the G2019S mutation that causes Parkinson’s disease. Mongolicain-A demonstrated good selectivity and specificity for the G2019S mutation in the target protein, making it a suitable candidate for leading the process towards drug discovery. Since guava has been previously reported to help manage neurodegenerative illnesses by lowering oxidative stress and neuro-inflammation, it’s a positive affirmation for tackling the debilitating Parkinsonism affecting millions of people worldwide.

Bacoside-A derived from *Bacopa monnieri is* already known to protect neurons against oxidative damage^21^and A-induced toxicity^22^. Molecular simulation and docking studies confirmed the above by displaying a high binding affinity toward the target protein. Bacoside A, on the other hand, was non-specific for the mutated ROCO 4 protein and other kinases, making it less efficient than Mongolicain-A.

But a vital obstruction to the treatment of any neurodegenerative disorder is the passage of drugs through the blood-brain barrier (BBB). The lack of proper cures and disease-modifying agents for brain disorders is partly due to the blood-brain barrier that blocks the distribution of potential drugs in the central nervous system. Mongolicain A has a molecular weight of approximately 1 kDa and is a non-BBB-permeable compound. However, recent scientific advances have introduced several physical and chemical methodologies to ease the delivery of target molecules with a high molecular weight into the brain by surpassing the blood-brain barrier. The Adsorptive Mediated Pathway (AMT) refers to an unspecific, clathrin-independent mode of transcytosis that allows the delivery of large molecules to the brain^23^. During this process, the anionic glycocalyx in the cell membrane binds to cationic molecules like albumin, allowing its delivery and entry to the brain. Thus, designing specific nano-particles encapsulating our drug followed by its binding to anionic albumin might be a possible intervention to improve its delivery through AMT. Besides macropinocytosis, which is also a non-specific mode of transcytosis (Smith & Gumbleton, 2006) *c*an be used for delivering our drug molecule. Further comprehensive experiments and studies are needed to understand the properties of the drug and finalize an efficient mode of administration.

Thus, results of this in-silico experiments identify Mongolicain-A as a potential LRRK2 (G2019S) inhibitor. Previous studies and experiments have already confirmed the potentiality of phytochemicals in maintaining vital body functions and exhibiting neuroprotective properties, and these studies partly support this hypothesis. But further in-vivo biological, chemical, and toxicity analysis of the compounds has to be performed to verify the above and their usage as a drug molecule in humans. This study would further provide light on options for treating Parkinson’s disease, which remains a great challenge in the medical field.

